# Social interaction and rate effects in models of musical synchronization

**DOI:** 10.1101/2022.02.24.481865

**Authors:** Valentin Bégel, Alexander P. Demos, Michelle Wang, Caroline Palmer

**Author notes:** Correspondence to: Prof. Caroline Palmer or Dr. Valentin Bégel, Dept of Psychology, McGill University, 1205 Dr. Penfield Ave. Montreal QC H3A 1B1 Canada.

## Abstract

Important individual differences are observed in people’s abilities to synchronize their body movements with regular auditory rhythms. We investigate whether synchronizing with a regular auditory cue is affected by each person’s spontaneous production rate (SPR) and by hearing a partner’s synchronization in a social context. Musically trained and untrained participants synchronized their tapping with an auditory cue presented at different rates (their own SPR or their partner’s SPR) and in a Solo or Joint (turn-taking) condition. Linear and nonlinear oscillator models were fit to participants’ mean asynchronies (signed timing differences between the cued onsets and taps). In Joint turn-taking, participants’ synchrony was increased when the auditory signal was cued at the participant’s own SPR, compared with their partner’s SPR; in contrast, synchronization did not differ across rates in the Solo condition. Asynchronies in the Joint task became larger as the difference between partners’ spontaneous rates increased; the increased asynchronies were driven by the faster partner who did not slow down to match the rate of their slower partner. Nonlinear delay-coupled models (with time delay, coupling strength, and intrinsic frequency) outperformed linear models (intrinsic frequency only) in accounting for tappers’ synchronization adjustments. The nonlinear model’s coupling value increased for musically trained participants, relative to untrained participants. Overall, these findings suggest that both intrinsic differences in partners’ spontaneous rates and the social turn-taking context contribute to the range of synchrony in the general population. Delay-coupled models are capable of capturing the wide range of individual differences in auditory-motor synchronization.

## Introduction

Many types of human behavior require individuals to coordinate the timing of their actions with external sounds. In social activities, including conversational speech (Wilson & Wilson, 2005) and ensemble music-making (Hadley, Novembre, Keller, & Pickering, 2015), coordination is often achieved by members taking turns with partners and anticipating their timing behavior, but little is known of how turn-taking affects individuals’ timing performance. Tapping experiments have shown that humans are able to synchronize their movement with regular auditory cues such as a metronome, but important individual differences are observed in this capacity (Dalla Bella et al., 2017; Repp, 2010; Scheurich, Zamm, & Palmer, 2018; Zamm, Wellman, & Palmer, 2016). Previous work suggests that individual differences in synchronization with auditory signals are influenced by individuals’ natural tapping or music performance rate (Scheurich et al., 2018; Zamm, Pfordresher, & Palmer, 2015), called the spontaneous production rate (SPR). From a dynamical systems perspective, the SPR is thought to represent the natural frequency of an underlying periodic oscillator that drives a person’s ability to synchronize with a partner (Zamm et al., 2016; Palmer et al., 2019) or with a regular auditory stimulus such as a metronome (Scheurich et al., 2018; Zamm et al., 2018). Turn-taking, a natural social context for making sound with a partner, is an interesting case for study as each person’s productions may be influenced by those of a partner. Few studies have addressed synchronization abilities in a turn-taking context. We investigated how turn-taking affects the synchronization abilities of individuals as they tap with a regular auditory cue in the presence of a partner (social turn-taking) or in the absence of a partner (solo turn-taking with a metronome). We applied nonlinear models from a dynamical systems perspective to interpret how turn-takers influenced each other.

Joint music performance offers an interesting trade-off between ecological validity and experimental control to study social aspects of turn-taking behavior (D’Ausilio, Novembre, Fadiga, & Keller, 2015; Hadley, Novembre, Keller, & Pickering, 2015). One question is how the presence of a partner influences individuals’ timing precision when they produce musical sequences in synchrony with a pacing cue such as an auditory metronome. Hearing a partner’s performance in a turn-taking context may help participants synchronize more accurately by imitating their partner. The presence of an adult model or tutor appears to reduce tapping variability in toddlers (Kirschner & Tomasello, 2009), but it is unclear whether such a social context would help adults to synchronize, given their broader experience. Turn-taking with a poor synchronizer may in fact hinder their performance. Previous research with adults showed that learning to produce rhythmic sequences is more challenging when taking turns with a partner than when learning to produce the same sequences alone (Zamm, Debener, & Sebanz, 2021).

A consistent observation in auditory-motor synchronization studies is that individuals tend to anticipate a regular auditory cue rather than lag behind it (Palmer & Demos, 2022; Repp, London, & Keller, 2011). Two theoretical frameworks have been developed to explain this anticipatory behavior, those of “weak anticipation” and “strong anticipation” (Dubois, 2003). The weak anticipation theory proposes that individuals maintain an internal model of the world that makes predictions about future events; anticipation results from enacting the internal model’s predictions (Clark, 1997; Clark & Grush, 1999). In contrast, the strong anticipation framework claims that synchronization can be explained by coupled dynamical systems, where a “driver” oscillator entrains a “driven” oscillator that anticipates based on time-delayed information about past behavior (Voss, 2000). One of the strengths of strong anticipation is that it not only accounts for anticipatory behavior in humans (Demos et al., 2019; Stepp 2009; Washburn et al., 2019), but also in physical systems that are not able to make mental predictions about the world (Stepp & Turvey, 2010).

Strong anticipation arises when a driven oscillator (such as a human tapping with an auditory metronome) compares instantaneous feedback from a driver oscillator (such as the metronome) with time-delayed feedback from itself, a process called “delay-coupling” (Ciszak et al., 2004; Stepp & Turvey, 2010). Anticipatory behavior was observed in unidirectional delay-coupled model simulations (Pyragas, 1992). In delay-coupled models, a chaotic driver system is coupled with a driven simple harmonic oscillator based on coupling strength and time-delay parameters (Stepp & Turvey, 2010). The driven oscillator receives instantaneous feedback from the driver oscillator and compares it with its own time-delayed feedback (Ciszak et al., 2004; Stepp & Turvey, 2010), whereas the driver oscillator receives no feedback. The delay-coupling approach has been extended to study mutual anticipation in pianist pairs who performed together (Demos, Layeghi, Wanderley, & Palmer, 2019). Demos et al. (2019) applied a bidirectional delay-coupling model to the asynchronies of the pianist pairs under different conditions in which the auditory feedback about the partner’s behavior was present or absent, forcing each pianist to become the driver or the driven partner. The delay-coupling model captured the partners’ asynchronies based on the coupling of two oscillators (representing the pianist pair) which received time-delayed feedback from each oscillator. Despite the large number of studies of human synchronization with an auditory cue, to our knowledge the unidirectional delay-coupled model has not been applied to turn-taking synchronization.

A dynamical system’s oscillation rate is often a control parameter that influences the system’s degree of synchrony with an external rhythm. The intrinsic frequency of each oscillator in a delay-coupled model corresponds to the spontaneous frequency (rate) at which it oscillates (Demos, Layeghi, Wanderley, & Palmer, 2019; Stepp & Turvey, 2010). Synchronization among different oscillations or individuals is more easily achieved and requires less coupling when the difference in their intrinsic frequencies is small. Models of human synchronization often estimate an intrinsic oscillator frequency from the spontaneous (uncued) rate at which a participant produces a familiar auditory sequence such as music (Palmer et al., 2019; Scheurich, Zamm, & Palmer, 2018; Zamm, Wellman & Palmer, 2018). Evidence to support spontaneous rates as representative of an intrinsic frequency is based on findings that both musicians and nonmusicians synchronize more accurately with an external stimulus when its rate is closer to their spontaneous rate (Scheurich, Zamm, & Palmer, 2018). In addition, musician partners with similar spontaneous rates achieve greater interpersonal synchrony in joint performance than those with dissimilar rates (Zamm, Wellman, & Palmer, 2016). Pianists display better synchronization accuracy when they synchronize with a recording of their own performance than with other performances (Keller, Knoblich & Repp, 2007), reflecting better temporal matching with sequences produced at the participant’s own spontaneous rate. Finally, individuals tend to drift closer to their spontaneous rates and show greatest temporal precision when performing at rates close to their spontaneous rates (Zamm, Wang, & Palmer, 2018). It is unknown, however, whether these effects of intrinsic frequencies on performance are influenced by the social context, such as that of turn-taking.

The range of success in interpersonal synchrony may depend on an interaction of the social context with the constraints introduced by intrinsic frequencies. Interestingly, people report increased social connection (Lumsden et al., 2014; Tarr, Launay, & Dunbar, 2016) and positive feelings (Hove & Risen, 2009) about a person with whom they engage in synchronous behavior. A possible explanation for the link between interpersonal synchrony and social interaction is that synchronization skills displayed by humans could be grounded in social interaction (e.g., Savage et al., 2021). Thus, large differences in the intrinsic frequencies of individuals who take turns synchronizing may yield reduced temporal coordination and reduced positive feelings and social connection with their partners, relative to partners with similar intrinsic frequencies.

This study assessed participants’ synchronization accuracy and perceived social interaction in a social turn-taking task. Musically trained and untrained partners synchronized their taps with auditory cues whose rates corresponded to the partners’ intrinsic frequencies. We hypothesized that synchronization accuracy would be higher when the cued rate was set to the participant’s own SPR instead of a partner’s SPR, and that the social context (Solo or Joint (turn-taking) performance) would influence the partners’ synchrony with the auditory cue. We then fit nonlinear and linear computational models to the synchronization behavior. To avoid the dangers of overfitting that can arise with complex nonlinear models (Hastie et al., 2017), we applied cross-validation methods, which consists of first fitting the model to a portion of the experimental data (the “Train” set) and then evaluating its fit to the other portion of the data (the “Test” set) while keeping model parameters fixed (Hastie et al., 2017; James et al., 2013). Cross-validation measures thus allow the comparison of multiple models whose parameters are optimized while minimizing overfitting. We divided our experimental data into a Train and a Test set, and we compared their model fits. To evaluate the goodness of model fits to each partner’s Test data, we compared them with model fits to data from other participants (“Surrogate” data). A significant difference between the model fits to the Test and Surrogate data would indicate that the models were able to capture individual differences between participants. Finally, we compared the participants’ ratings of social interaction with their partner to the model parameters, amount of musical training, and the behavioral asynchronies.

## Methods

### Participants

Twenty-four participants were recruited. Half of the participants, referred to as Musicians (7 women; age range: 19–32 years, *M* = 22.67, *SD* = 4.42), had at least six years of private instrumental musical training (range: 6–16 years, *M* = 9.5, *SD* = 3.15) and were actively playing their instrument at the time of the study. The other half, referred to as Nonmusicians (8 women; age range: 19–30 years, *M* = 22.83, *SD* = 3.24), had less than or equal to two years of private instrumental musical training (range: 0–2 years, *M* = .42, *SD* = 0.76) and were not actively playing an instrument at the time of the study. All participants were right-handed. Participants who reported a neurological condition were not included in the study. An audiometry screening was administered with a Maico MA40 audiometer. All participants had normal hearing in the frequency range of stimuli used in the experiment (< 30 dB HL for single tones in the 125–750 Hz pitch range). Four participants (all musicians) who were tested during the COVID-19 pandemic did not perform the audiometry screening in order to minimize risks of COVID-19 transmission, and the decision to end data collection after n=24 participants was made due to COVID-19 restrictions. Participants received a small monetary compensation for their time, and the study was approved by the McGill University Research Ethics Board.

### Equipment and Stimuli

Participants tapped musical melodies on a force-sensitive pad controlled with Arduino and connected to a Linux computer (Dell T3600 running Fedora 16) by a MIDI cable. They heard the melodies they tapped through headphones (AKG K240 Studio) in a marimba timbre (GM2, patch #13, bank #0) produced with a Roland Studio Canvas SD-50 tone generator. Metronome beats were presented with a high-pitched woodblock timbre (GM2, patch #116, bank #0). The timing of the sounds was controlled with the FTAP program (Finney, 2001), which ensured a negligible time delay from the start of the tap on the pad to the start of the sound (less than 1.0 millisecond; Scheurich, Zamm, & Palmer, 2018, Supplemental materials).

An 8-note melody corresponding to an ascending C major scale, starting at C4 and ending at C5, was used for each partner’s feedback in all experimental conditions. This melody was chosen for its familiarity regardless of participants’ musical background and because it is natural to hear and perform the scale with an isochronous rhythm. Each partner’s feedback was sounded with the same timbre in the same pitch range in order to present participants with equivalent acoustic cues.

### Design

The experiment included a Group factor of musical expertise (Musician / Nonmusician) which was the only between-subject variable; Musicians were randomly paired with Musicians, and Nonmusicians were randomly paired with Nonmusicians. The experiment consisted of three tasks, performed in the following order: a Spontaneous Rate (SPR) task, a Solo synchronization task (turn-taking with a metronome), and a Joint synchronization task (turn-taking with a partner). The tasks are described further below. Individuals’ mean interonset intervals (IOIs) from the Spontaneous Rate task were used to determine the auditory cued Rate variable in both Solo and Joint tasks; each partner pair performed the Solo and Joint tasks at both their Self-cued Rate and Partner-cued Rate. The order of Rate conditions was fixed to Self followed by Partner in the Solo task, and was assigned randomly in the Joint task. Each of the Solo and Joint tasks presented at each of 2 Rates contained 3 trials, yielding a total of 2 × 2 × 3 trials or 12 trials. Thus, the synchronization tasks followed a mixed design with the between-subject factor of Group (Musician and Nonmusician) and the within-subject factors of Cued Rate (Self SPR, Partner’s SPR) and Task (Solo, Joint).

### Procedure

Participants were scheduled for the experiment at the same time as a randomly paired partner. Participants gave informed consent prior to or upon arrival at the lab. Each partner first performed an audiometry screening, the SPR task, and the Solo synchronization task in different rooms. Then, the partners performed the Joint synchronization task together in the same room.

Participants were first presented with the SPR tapping task. They were told to tap on the force-sensitive pad using the index finger of their dominant hand and that each tap would produce the next tone in the familiar melody, heard over headphones. They first completed two example trials to become comfortable with the task, tapping the 8-note melody twice without stopping between repetitions. The experimental trials consisted of tapping the melody 4 1/2 times in the absence of any rhythmic pacing cue. They were instructed to tap at a steady and comfortable rate, without stopping between repetitions until the taps generated no more auditory feedback. Participants performed one practice trial and three experimental trials. Any trial with obvious rhythmic mistakes was replaced with an additional trial. After the SPR task, participants were asked to complete a musical background questionnaire (except for the last four participants, who had previously completed the questionnaire online). After this task, the participants’ SPRs were calculated to determine the metronome rates in the synchronization tasks.

Next, participants were presented with the Solo synchronization task. They first performed a practice trial in which they tapped the 8-note melody twice, without stopping between repetitions, while synchronizing with a high-pitched auditory cue that was sounded every 500 ms. Then participants were instructed to alternate between synchronized tapping for 8 cued beats and waiting for 8 cued beats (see Figure 1A top for a sample trial). Each trial lasted 4 1/2 repetitions, until no more auditory feedback was heard. Then the experimental trials began in which the auditory cue was set to the Participants’ Rate from the SPR task. Participants performed a practice trial followed by three experimental trials.They then repeated this procedure of one practice trial followed by 3 experimental trials with the metronome cue set to the rate at which their partner had performed the SPR task (Partner’s SPR).

**Figure 1.**
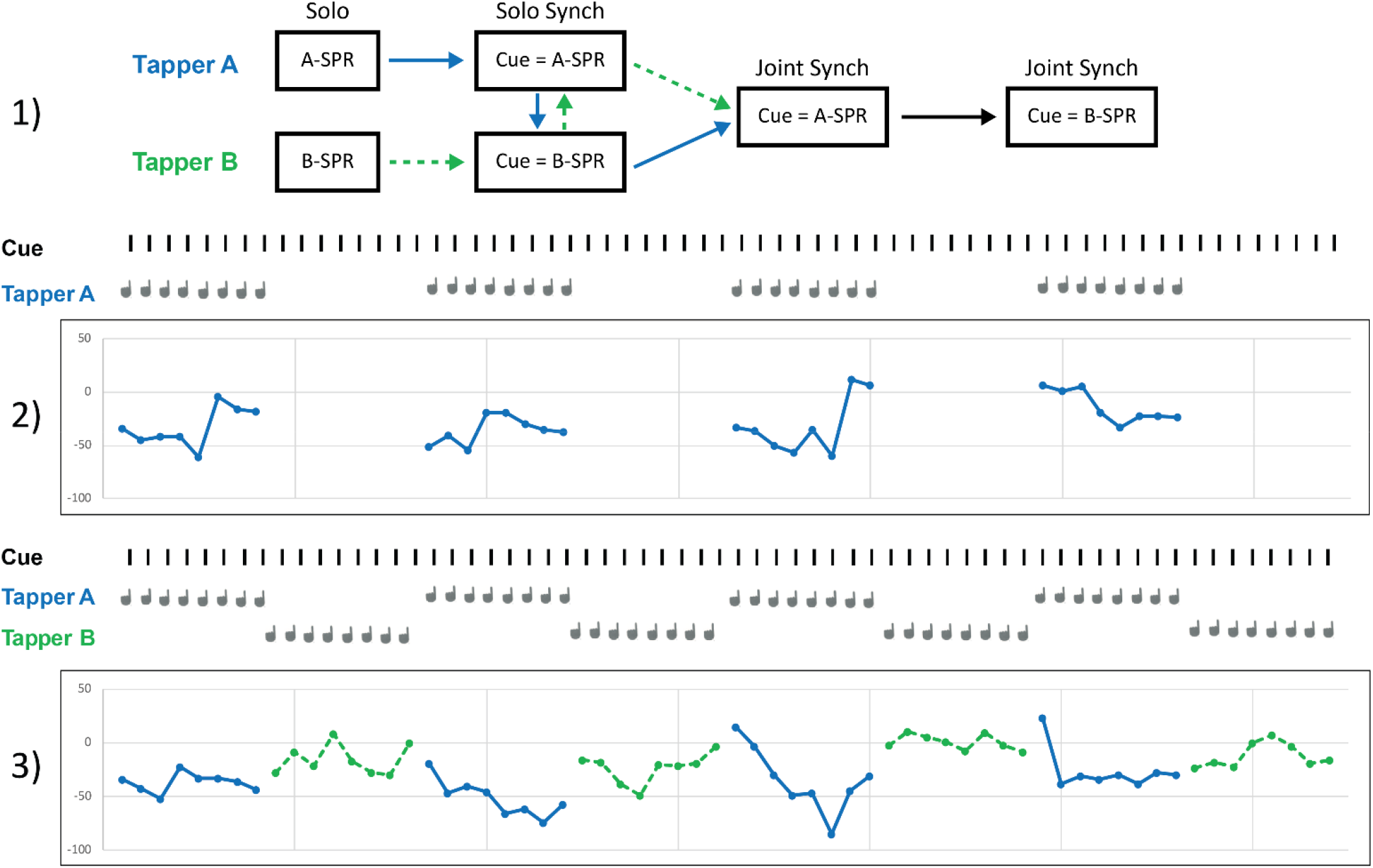
Schematic representation of the experimental procedure. 1) Each partner completes the Solo SPR task, followed by the Solo Synchronization task first with the auditory cue set to their own SPR and then with the auditory cue set to their partner’s SPR. The two partners then complete the Joint (Turn-taking) Synchronization task first with the auditory cue set to one partner’s SPR and then to the other partner’s SPR. 2) Sample trial from one participant (Tapper A) in the Solo synchronization task with the auditory cue set to Tapper A’s SPR (545 ms ITI). 3) Sample trial from two participants in the Joint synchronization task with the auditory cue set to Tapper A’s SPR (545 ms ITI)

Finally, the partners performed the Joint (turn-taking) synchronization task in the same room as their partner. They faced each other at two separate tables, each with its own force-sensitive pad setup, and a screen was placed between them so that they could only see their partner’s head and shoulders, to avoid influences of finger movements. Both participants changed their physical locations from the Solo task, so that the Joint task was equally novel for the two participants. Each participant was instructed in the Joint turn-taking task to alternate between synchronized tapping for 8 metronome beats and waiting while their partner tapped for 8 metronome beats (see Figure 1B bottom for a sample trial). Each partner heard the metronome cue for 8 beats at the beginning of the trials, as well as their own tapping feedback and their partner’s tapping feedback. Each trial lasted 4 1/2 melody repetitions or 4 sets of 8 taps per partner (the same amount of taps per partner as in the Solo synchronization condition), until no more auditory feedback was sounded. The participants first performed a practice trial in which the metronome was cued at 500 ms. They then performed a practice trial with the metronome cue set to one partner’s SPR, followed by three experimental trials cued at the same rate. The first partner to tap was always the person whose SPR determined the metronome cue on that trial. The Joint synchronization task was then repeated with the metronome cue set to the other partner’s SPR, and that partner began the tapping.

At the end of the experiment, participants were asked to complete a questionnaire with three questions presented on a 7-point Likert scale: how pleasant was the task (pleasantness), how close was their relationship with their partner (relationship), and how successful they thought their synchronization was (synchronization). Four participants who were tested during the COVID-19 pandemic did not complete the questionnaire. The entire experiment lasted approximately 60 minutes and participants received a small fee for their participation.

### Data Analysis

Each participant’s rate in the SPR task was calculated as the mean intertap interval (ITI, in ms) across the middle two repetitions (most stable sections) of each experimental trial of the SPR task, similar to other studies (Zamm, Wellman, & Palmer, 2016; Zamm, Wang, & Palmer, 2018; Palmer et al., 2019). The Coefficient of Variation (CV) of the taps was calculated as the SD of the same ITIs divided by the mean ITI. Each tap in the Solo and Joint synchronization trials was matched with an auditory cued beat following a nearest-neighbour approach. Signed asynchrony values were computed for Solo and Joint synchronization trials as tap onset time minus cue onset time in ms. Hence, a negative asynchrony value indicated that the tapper anticipated (tapped before) the cue, while a positive value indicated that the tapper lagged behind the cue. Variability indices were also calculated for the synchronization tasks based on the CV of ITIs and the SD of the signed asynchronies.

### Delay-coupled model

The unidirectional delay-coupled model in which a driven oscillator couples with a driver oscillator can be described by the equations below (Stepp & Turvey, 2010).

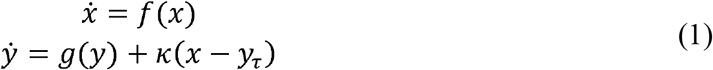

In Equation 1, *x* represents the driver oscillator (the auditory cue) and *y* represents the driven oscillator (the participant). The functions *f (x)* and *g (y)* describe the behavior of the oscillators which depends on their intrinsic frequencies. The driver oscillator *x* does not receive any feedback, whereas the driven oscillator y receives feedback from x that is compared to its delayed self-feedback *y*_*τ*_ *= y (t − τ)*, where *τ* is a time delay parameter. The parameter *κ* is the coupling strength between the driver and driven oscillator.

Equation 2 defines the delay-coupled model in which constant values are assumed for the intrinsic frequencies of *f (x)*and *g (y)* (Demos, Layeghi, Wanderley, & Palmer, 2019):

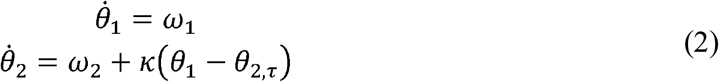

*θ* _1_ and *θ*_2_ denote the phase of the driver metronome and the participant, respectively, and *ω*_1_ and *ω*_2_ denote their intrinsic frequencies. The driver’s intrinsic frequency parameter was fixed at the auditory metronome rate. The coupling term κ influences the difference in relative phase between the oscillators. The value of the time delay parameter *τ* is assumed to be a constant value, thought to reflect neural transmission delays (Machado & Matias, 2020; Roman et al., 2019). The time delay affects the degree of anticipation of the driven system relative to the driver. Thus, the model was fit on two parameters, *κ* and *ω*_2_. The intrinsic frequency of the driven oscillator (*ω*_2_) was modeled as *ω*_*diff*_, the signed difference between the intrinsic frequency and the cued metronome rate (*ω*_*diff*_ = *ω*_2_ -*ω*_1_).

The delay-coupled model was fit to the mean asynchrony values of each trial for each participant. The model-fitting procedure consisted of two stages: in the first stage, a global parameter search was conducted using a genetic algorithm. In the second stage, the previously obtained parameters were passed to a local search algorithm (constrained nonlinear multivariate function). The error function was a weighted sum-of-squared-errors function where the first serial position was weighted by 4 and the eighth (last) position was weighted by 2, similar to what Demos et al. (2019) used to capture the curvilinear relationship of the asynchrony patterns following perturbations in the auditory feedback heard right before each synchronization sequence. The Solo and Joint experimental tasks can also be viewed as introducing perturbations, since participants heard alternate patterns of auditory feedback as they waited 8 beats between their turns tapping. The model was fit 10 times on the averaged series of 8 asynchronies for each condition and for each participant, and the model whose parameters generated the best (lowest) root mean squared error (RMSE) was chosen for subsequent analysis.

The delay-coupled model was first fit with the driven oscillator’s three parameters allowed to vary. The driven oscillator’s intrinsic frequency parameter *ω*_2_ was allowed to vary within a range of ± 300 relative to the value of the *ω*_1_ parameter (cued metronome rate). The coupling strength parameter was allowed to vary from 0 (no coupling) to 50 ms. The *τ* parameter was allowed to vary with a range of 0 (no time delay) to 50 ms.

Following these initial model fits, we fit the model again, this time using a fixed time delay *τ*, as in other studies (Machado & Matias, 2020; Roman et al., 2019), set to the median *τ* value (10.15 ms) from the initial model fits. The range of the *ω*_*diff*_ parameter was again allowed to vary within ± 300 of the value of the *ω*_1_ parameter. Two of the participant pairs had differences in the partners’ SPR values larger than 300ms (the maximum range allowed for *ω*_2_ to change); therefore the range of the *ω*_2_ parameter was increased to ± 350 for these 2 pairs to reduce the number of boundary hits.

### Linear model

A linear model was fit to the asynchrony data the same way as for the delay-coupled model, with the only difference being that both the *κ* and the *τ* parameters were fixed at zero. We compared the delay-coupled model with the simpler, linear model defined by Equation 3.

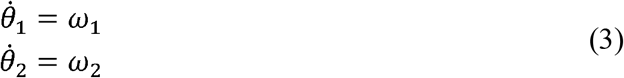

As with the delay-coupled model, the *ω*_1_ parameter was fixed to the metronome’s rate. Only the *ω*_2_ parameter was allowed to vary during model fitting within ± 300 of the value of the *ω*_1_ parameter (± 350 for the 2 pairs with differences in the partners’ SPR values larger than 300ms).

### Cross-validation and surrogate analysis

The behavioral asynchrony measures were divided into a Train set and a Test set for the model comparisons. Each trial from the Solo or Joint synchronization tasks consisted of four repetitions of the eight-note melody. The first and third repetitions within each participant’s trial were averaged together to form a single series of eight asynchronies which were assigned to either the Train or the Test set (in equal amounts), and the second and fourth repetitions were averaged together and assigned to the other set (Train or Test). The repetitions were paired together this way (1-3 and 2-4) to account for potential effects of serial position within the trial. Optimal values for the model parameters were then determined for each trial based on the Train dataset, using the same model fitting procedure as previously described. Both the delay-coupled model and the linear model were fit to the Train dataset. For the delay-coupled model, two trials were removed because the parameter values reached ceiling, indicating a poor fit (within 0.01 of the given upper bound). None of the linear model trials reached ceiling in model fits.

We then compared the *predicted* asynchronies based on the model fits to the Train datasets with the *observed* asynchronies in the Test dataset, using z-transformed Pearson correlation values as a measure of goodness-of-fit. To generate a chance estimate for the Train-Test correlations, a surrogate reference was created for the Test dataset by comparing the *predicted* asynchronies for a given partner with *observed* data from partners in other pairs, called “Surrogates”. For each trial of a partner, a Surrogate comparison was created by taking one trial from one partner in each of the other pairs of participants, using the same trial conditions (same Task, Cued Rate, trial number, and repetitions) as in the Test data. Each of the Surrogate trials was then correlated with the model’s predicted Train dataset asynchronies for each partner, and the correlation coefficients were averaged to obtain a single value that could be compared to the obtained Train-Test correlations. Since the Surrogate trials were not taken from the actual partner as was used in the Train/Test correlations, we expected the predicted asynchronies to not match the Surrogate data well. Therefore, the Train-Test correlation coefficients based on the Surrogate data served as chance estimates for the Test data: similar Train-Test and Train-Surrogate correlations would suggest that the models were not able to generalize to the Test set.

### Statistical Analyses

All analyses were run in R Statistical Software (v4.1-0; R Core Team, 2021). Tests of the behavioral measures included analyses of variance (ANOVA; Afex v1.0-1; Singmann et al., 2021) that addressed Group differences (Musician, Nonmusician), Cued Rate (Self or Partner), and Task (Solo, Joint) on SPR (IOI, CV) and Synchronization values (Asynchronies, CV of IOI, SD) with individuals coded as the random variable. Linear contrasts were run with the Emmeans package (1.7-2; Lenth, 2022) and p-values were corrected within family when there were more than two tests using a Tukey correction. Wilcoxon tests were applied to compare the social interaction responses of participants in “Uncoupled” pairs (i.e., pairs in which at least one participant showed no coupling in all three trials within at least one of the four experimental conditions) with participants in “Coupled” pairs. One-tailed tests were applied to the social interaction ratings to test the hypotheses that partners in the Uncoupled pairs would rate pleasantness, relationship, and synchronization as lower than participants in the Coupled pairs.

Mixed models were next applied to the observed asynchronies to further examine individual and pairwise differences in the partners’ data using the lme4 package in R (v1.1-8; Bates et al., 2015) and we report Type III F-tests with Saitherwaith degrees of freedom using the LmerTest package (v3.1-3; Kuznetsova, Brockhoff, & Christensen, 2017). Both the individual participants and the pair were included as random variables. The following variables were dummy coded as fixed effects: Faster or Slower Cued Rate and Task (Solo or Joint), Musical training (years of individual instruction on a musical instrument), and the Rate Difference (absolute difference between the tapper’s SPR and the rate of the metronome). The Pair’s slope of the Rate difference was entered as a random effect. The Subject’s main effect and interaction of Task and Cued Rate were entered as random effects.

The model fits and cross-validation analyses were first analyzed with ANOVAs applied to the RMSE values by Model (Delay-coupled, Linear) and Set (Train, Test, Surrogate). Simple Pearson correlations (Fisher’s *r*-to-*z* transform) between observed and predicted asynchronies were analyzed in ANOVAs by Cued Rate (Self SPR, Partner’s SPR), Task (Solo, Joint), Model (Delay-coupled, Linear) and Set (Train, Test, Surrogate). Finally, the model’s estimated parameters (*κ* and *ω*_2_ values) for each participant were analyzed by Musicianship, Cued Rate (Self SPR, Partner’s SPR) and Task (Solo, Joint).

Correlations were conducted between participants’ answers to the social interaction questions, the mean and SD of asynchronies, and the estimated model parameters (*κ* and *ω*_*diff*_). P-values were corrected for multiple comparisons using Bonferroni correction.

## Results

### Spontaneous Production Rates

We first examined the obtained SPR values (mean ITI per trial) across participants and assessed whether they differed between musicians and nonmusicians. Figure 2A shows the SPR distribution across participants, ordered from fastest to slowest. Figure 2B shows SPRs for each pair, ordered from the smallest pairwise difference in partners’ SPRs to the largest difference. A one-way ANOVA by Group (Musician, Nonmusician) revealed no main effect of musical training on mean SPR, *F*(1, 22) = 0.10, *p* = .76,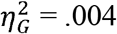; the mean SPR for musicians (484 ms, *SD* = 132.41) and nonmusicians (504 ms, *SD* = 168.36) did not differ significantly. Musicians were less variable in their SPR performance, as indicated by their lower CV of ITI (Musician, *M* = .034, *SD* = .009, Nonmusician, *M* = .60, *SD* = .029, *F*(1, 22) = 9.01, *p* < .001, 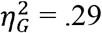).

**Figure 2.**
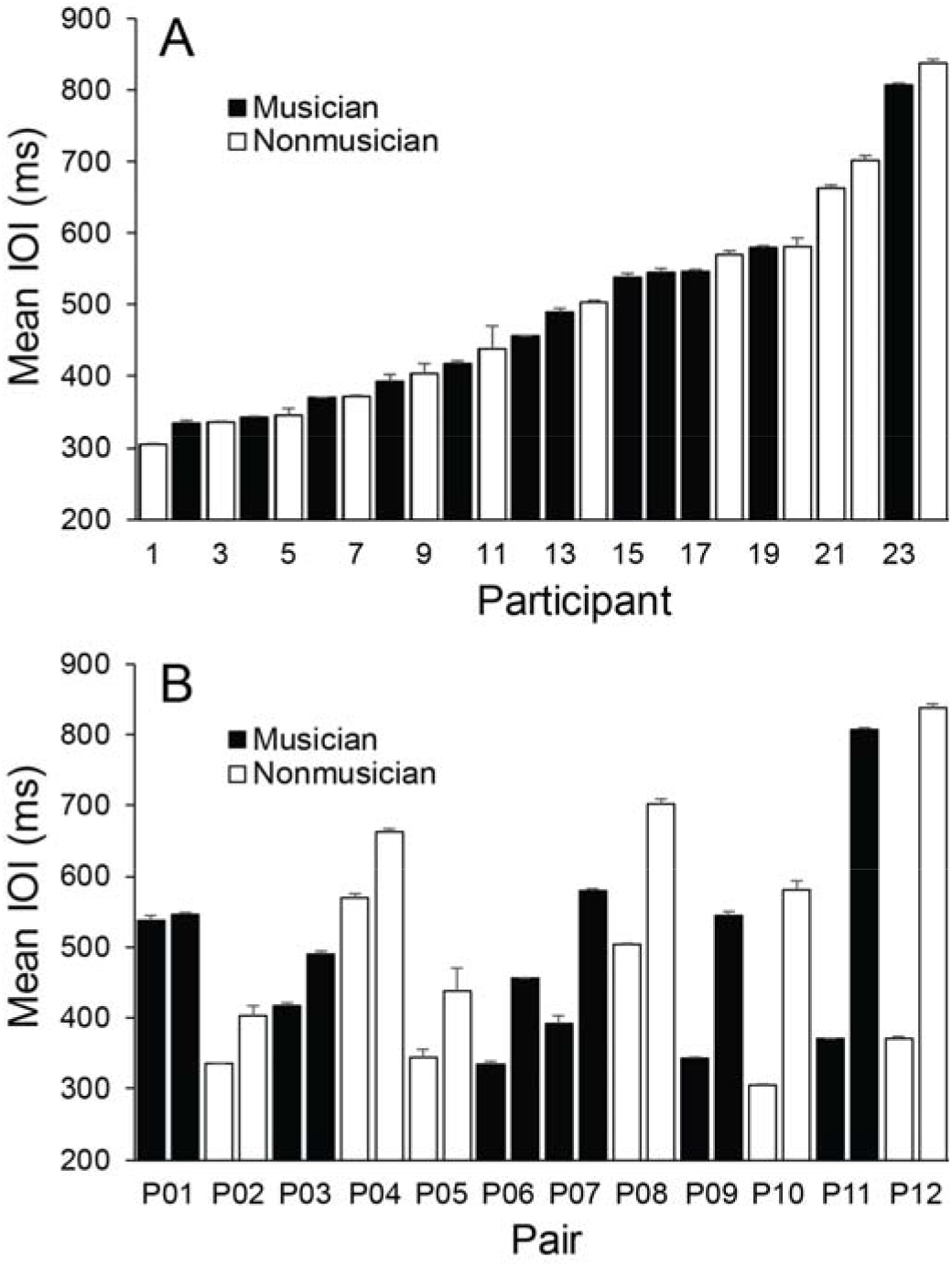
Mean individual Spontaneous Production Rates. Error bars indicate standard errors of the mean. A: Participants ordered by fastest SPR to slowest. B: Pairs ordered by smallest difference between partners’ SPRs to the largest difference.

### Synchronization accuracy

Figure 3 shows the participants’ mean asynchronies with the auditory cue from the synchronization tasks for each Cued Rate (Self SPR, Partner’s SPR) and Task (Solo and Joint). A three-way ANOVA on mean asynchronies by Group, Cued Rate and Task indicated a significant interaction between Cued Rate and Task, *F*(1, 22) = 6.68, *p* = .017, 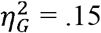. Participants’ mean asynchronies when tapping at their Partner’s Cued Rate were larger in the Joint task than in the Solo task, *t*(22) = 2.60, *p* = .016, *d* = 0.80. In contrast, the asynchronies in the Self Cued Rate condition remained small across Joint and Solo tasks, *t*(22) = 0.70, *p* = .489, *d* = 0.25. Participants’ mean asynchronies in the Joint task were marginally larger when tapping at their partner’s Cued Rate than at their own Self Cued rate, t(22) = 1.75, p = 0.094, d = 1.06. There was no difference between Cued rates in the Solo task, t(22) = 0.16, p = 0.987, d = 0.008. There were no main effects, or other interactions, of Group, Cued Rate or Task. Thus, the most difficult condition - that generated the largest asynchronies - was when the participant performed the Joint turn-taking task with the auditory cue set to the partner’s rate.

**Figure 3.**
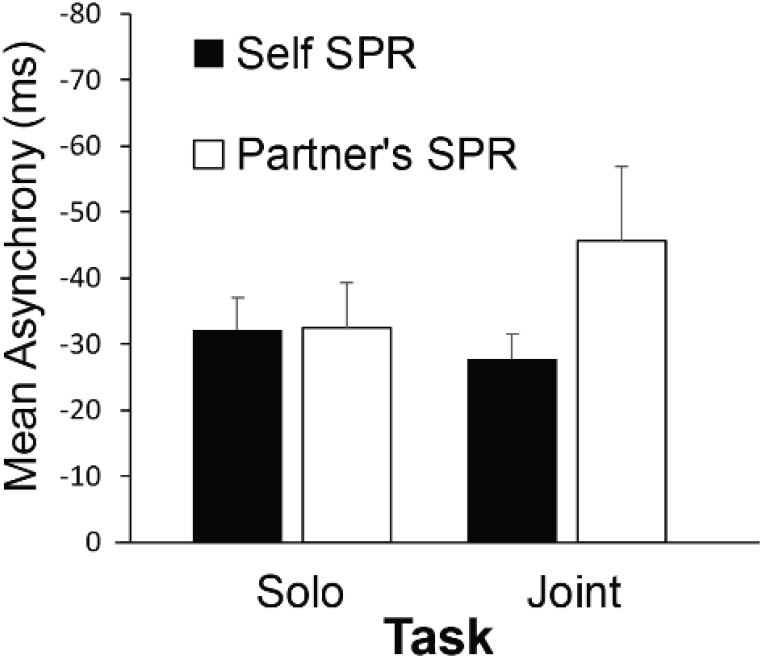
Mean asynchrony (tap onset minus auditory cue onset) by Task and Cued Rate. Error bars indicate standard error of the mean.

Next we compared effects of Group, Cued Rate and Task on CV of ITI measures and SD of asynchronies. Musicians’ CV of ITI (*M* = .090, *SD* = .044) were smaller than Nonmusicians’ (*M* = .317, *SD* = .082), *F*(1, 22) = 6.97, *p* = .015, 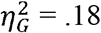, indicating lower variability in their tapping performance. Similarly, musicians’ SD of asynchronies (*M* = 25.61, *SD* = 28.32) were lower than Nonmusicians’ (*M* = 45.27, *SD* = 25.35), *F*(1, 22) = 6.26, *p* < .02, 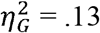. There were no other significant main effects or interactions.

Some pairs of partners had similar SPR values, whereas other pairs had larger differences in SPRs. To test the effects of this difference, a mixed model was applied to mean asynchronies by Faster or Slower Cued Rate, Task, Musical training (years of private instruction), and the absolute difference between the tapper’s SPR and the cued rate, called Rate Difference. A significant interaction was found between Task and Rate Difference, *F*(1, 27.04) = 6.78, *p* = .015. Figure 4 shows the asynchronies relative to the Rate Difference in the Solo and Joint conditions. Asynchronies became larger as the Rate Difference between the tapper’s SPR and the cued rate increased in the Joint condition; asynchronies remained similar across the Rate Differences in the Solo condition. Thus, performing at a partner’s SPR rate negatively affected participants’ synchronization in the Joint condition as the partner’s rate differed nor from their own rate. Also shown in Figure 4 is the mean asynchrony when tappers performed at their own rate (x-axis values = 0). There was no difference at the 0 point between the Solo and Joint tasks, *t*(48.88) = 0.36, *p* = .71, confirming that partners could synchronize accurately when they were cued at their own rate in Solo and Joint tasks.

**Figure 4.**
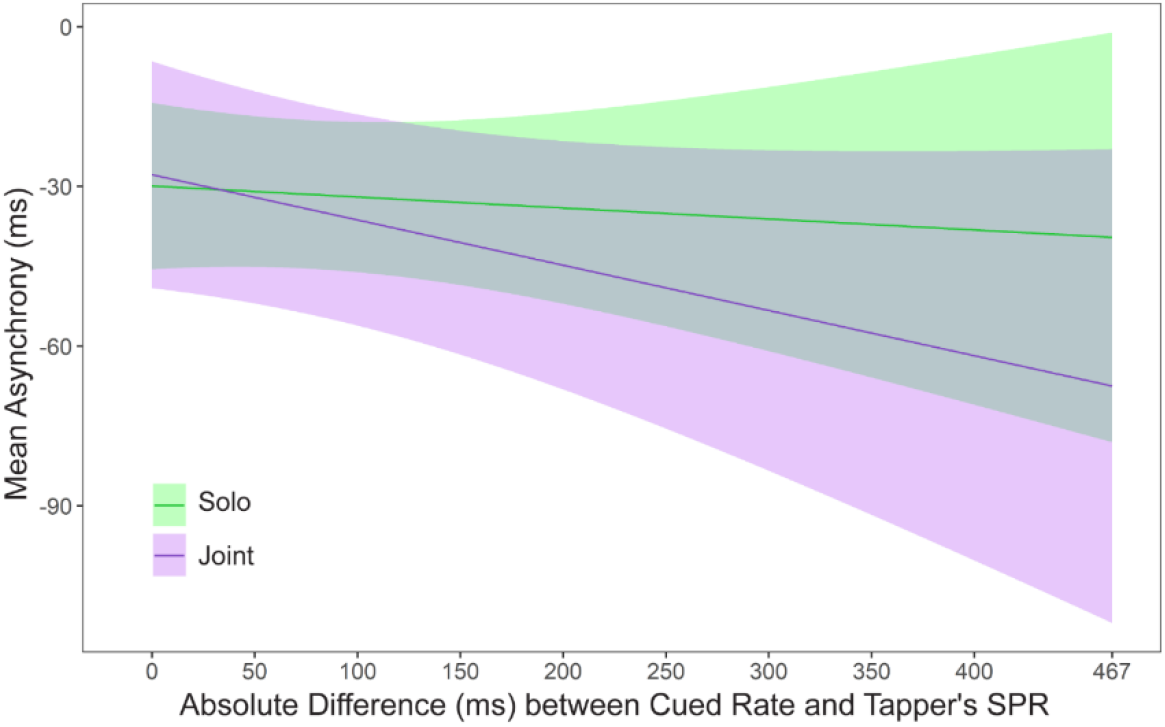
Asynchronies predicted by the mixed model: Interaction between SPR Difference and Task. Zero on the x axis represents the condition in which the metronome is cued at the tapper’s own rate.

An interaction was also found between Slower and Faster Rates and the Rate Difference, *F*(1, 45.35) = 7.54, *p* = .009. Figure 5 shows the predicted asynchronies by Slower/Faster rate and the Rate Difference. As seen in Figure 5, the faster partner in each pair was responsible for the larger asynchronies observed as the Rate Difference increased. Thus, the faster partner could not slow down to match the slower partner’s rate, whereas the slower partner could speed up to match the faster partner’s rate. There was no difference at the 0 point between the slow and fast Cued Rate, *t*(62.57) = 0.48, *p* = .63, confirming that partners could synchronize accurately when they were cued at their own rate in Slow and Fast tasks.

**Figure 5.**
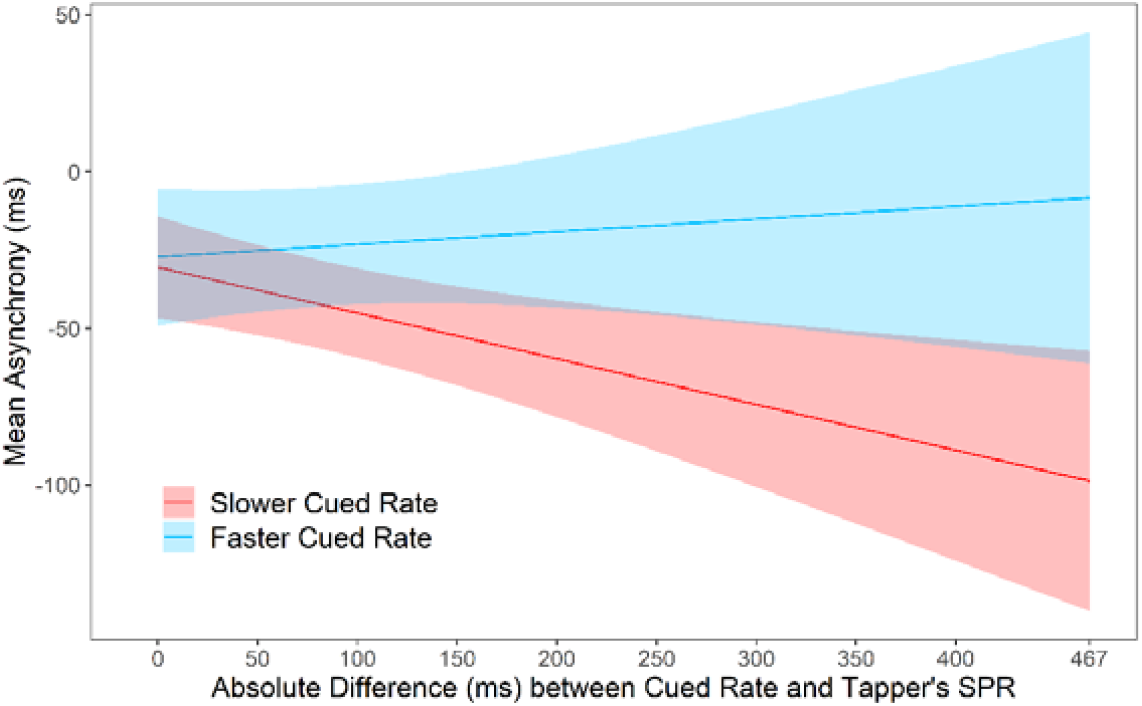
Asynchronies predicted by the mixed model: Interaction between Slower/Faster Cued Rate and the absolute difference between the tapper’s SPR and the rate of the metronome. Zero on the x axis represents the condition in which the metronome is cued at the tapper’s own rate.

#### Summary of behavioral results

Although the mean SPR for musicians and nonmusicians did not differ, musicians were less variable in both their SPR performances and their synchronization accuracy. Participants’ mean asynchronies in the Joint turn-taking task were larger when the auditory cue was set to their partner’s SPR than to their own SPR. Asynchronies increased in the Joint condition as the difference between partners’ SPRs increased. Those larger asynchronies were driven by the faster person in the pair, who could not slow down to match the rate of the slower partner.

### Model fits: Cross-validation and surrogate analysis

We identified participants who did not couple with the metronome by applying the linear model in which *k* = 0 to the asynchrony measures in the Solo and Joint synchronization conditions. If the RMSE of the linear model was less than (better) or equal to the RMSE of the delay-coupled model, then the participant was deemed as not having successfully coupled to the metronome during that trial. Figure 6 shows an example of a participant whose trial was better fit by the linear (uncoupled) model and one whose trial was better fit by the delay-coupled model. There were 89 trials out of a total of 288 (3 trials × 2 cued rates × 2 tasks × 24 participants) in which the RMSE of the linear model outperformed the delay-coupled model and no coupling was observed. The majority (94%) of these trials came from 7 participants, who were unable to adapt to the cued rate. These seven participants were the only participants to show no coupling in all trials of at least one of the four experimental conditions. The following statistical analyses were performed on data from n=163 trials for each model.

**Figure 6.**
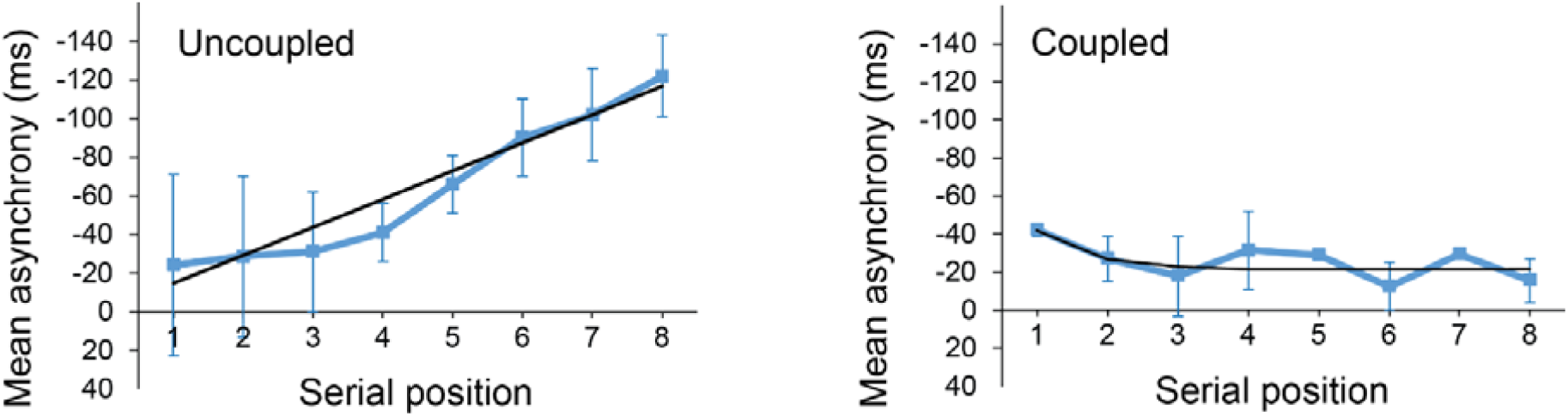
Mean asynchronies and model fits for a trial better fit by the linear (uncoupled) model (left) and a trial better fit by the delay-coupled model (right). The black line represents the model predicted asynchronies.

The delay-coupled and linear models were next fitted to the tappers’ asynchronies (*n* = 17). Figure 7 shows mean RMSE values obtained for the best-fitting delay-coupled model and linear model for each of the three datasets: Train, Test, and Surrogate, averaged across the four experimental conditions. A two-way ANOVA by Model (Delay-coupled, Linear) and Set (Train, Test, Surrogate) showed a significant main effect of Set, *F*(1.34, 21.48) = 69.01, *p* < .001, 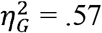. As expected, RMSEs for the Train set (*M* = 15.38, *SD* = 19.21) were smaller (better) than those for the Test set (*M* = 27.15, *SD* = 23.88), *t*(16) = −4.80, *p* < .001, *d* = 0.60, which in turn were smaller than those for the Surrogate set (*M* = 59.58, *SD* = 32.24), *t*(16) = −6.56, *p* < .0001, *d* = 1.65. There was also a significant main effect of Model, *F*(1, 16) = 6.74, *p* = .02,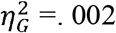. The delay-coupled model had smaller RMSEs (*M* = 33.36, *SD* = 31.17) than the linear model (*M* = 34.72, *SD* = 32.33). There was also a significant interaction between Set and Model, *F*(1.30, 20.83) = 4.00, *p* = .03, 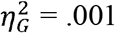): in the Train set, RMSEs were smaller (better) for the delay-coupled model (*M* = 13.97, *SD* = 18.66) than for the linear model (*M* = 16.80, *SD* = 19.79), *t*(16) = −5.33, *p* < .0001, *d* = 1.22.

**Figure 7.**
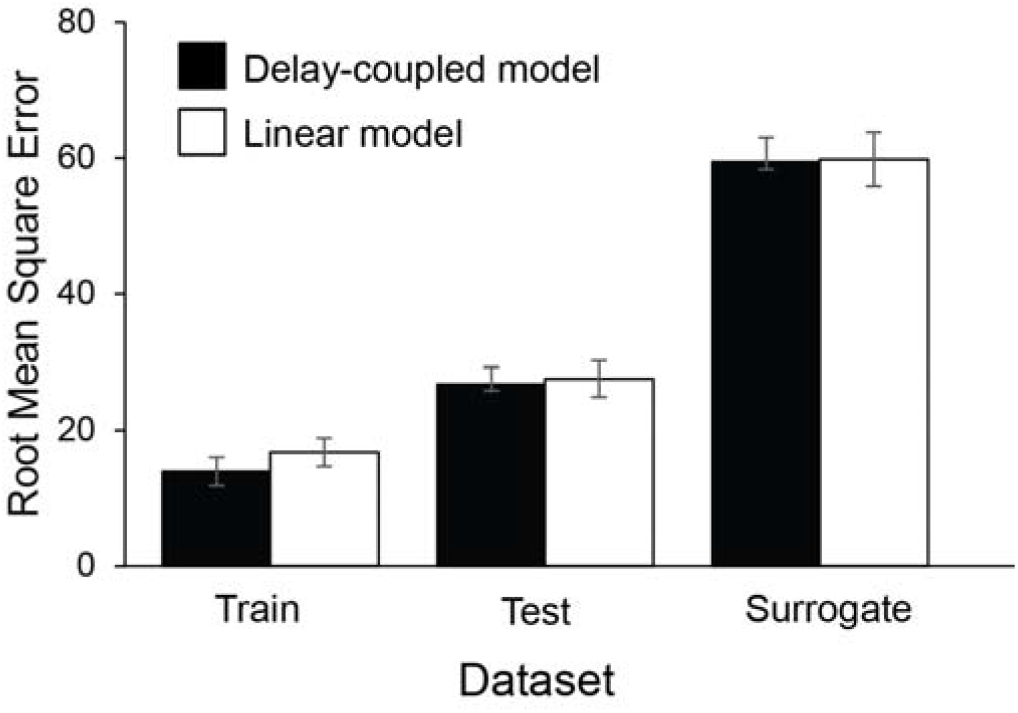
Root Mean Squared Error computed between the model’s predicted asynchronies and the participants’ observed asynchronies, by dataset and Model type.

A four-way ANOVA by Cued Rate (Self SPR, Partner’s SPR), Task (Solo, Joint), Model (Delay-coupled, Linear) and Set (Train, Test, Surrogate) on the z-transformed Pearson correlation coefficients showed a significant main effect of Model, *F*(1, 16) = 121.84, *p* < .001,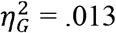. Correlation coefficients were larger (indicating better model fits) for the delay-coupled model (*M* = 0.65, *SD* = 0.66) than the linear model (*M* = 0.53, *SD* = 0.61). The ANOVA also revealed a significant main effect of Task, *F*(1, 16) = 8.74, *p* = .009, 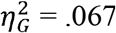, with larger correlation coefficients for the Solo task (*M* = 0.73, *SD* = 0.60) than the Joint task (*M* = 0.45, *SD* = 0.64), suggesting that the presence of a partner reduced the correspondence between observed and predicted values. There was also a significant main effect of Set, *F*(2, 32) = 38.20, *p* < .001, 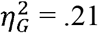: correlation coefficients for the Train set (*M* = 0.93, *SD* = 0.56) were larger than those for the Test set (*M* = 0.58, *SD* = 0.71), *t*(16) = 5.50, *p* = .0001, *d* = 0.46, which in turn were larger than those for the Surrogate set (*M* = 0.25, *SD* = 0.41), *t*(16) = 3.32, *p* = .011, *d* = 0.43.

The analysis indicated two significant interactions: one was between Cued Rate and Model, *F*(1, 16) = 6.76, *p* = .02, 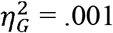 (Figure 8A). The difference between the two model fits was significantly larger in the delay-coupled model (*M* = 0.69, *SD* = 0.67) than in the linear model (*M* = 0.53, *SD* = 0.60) when the cued rate was the participant’s own SPR, *t*(16) = 8.69, *p* < .0001, *d* = 1.19, and when the cued rate was the partner’s SPR (delay-coupled model, *M* = .61, *SD* = 0.65; linear model, *M* = 0.52, *SD* = 0.63), *t*(16) = 5.50, *p* < .0001, *d* = 0.68, but the difference between the two models was larger when the cued rate was the participant’s own SPR. There was also a significant interaction between Model and Set, *F*(2, 32) = 43.76, *p* < .001, 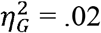 (Figure 8B). For the Train dataset only, correlation coefficients for the delay-coupled model (*M* = 1.09, *SD* = 0.54) were also significantly higher than for the linear model (*M* = 0.77, SD = 0.55), *t*(16) = 11.87, *p* < .0001, *d* = 0.42. For the Test dataset only, correlation coefficients for the delay-coupled model (*M* = 0.61, *SD* = 0.69) were marginally higher than for the linear model (*M* = 0.56, *SD* = 0.72), *t*(16) = 1.89, *p* = .076, *d* = 0.07. For the Surrogate dataset only, the correlation coefficients for the delay-coupled model (*M* = 0.25, *SD* = 0.42) and the linear model (*M* = 0.26, *SD* = 0.41) were not significantly different, *t*(16) = 0.51, *p* = .61, *d* = 0.01. There were no other significant main effects or interactions.

**Figure 8.**
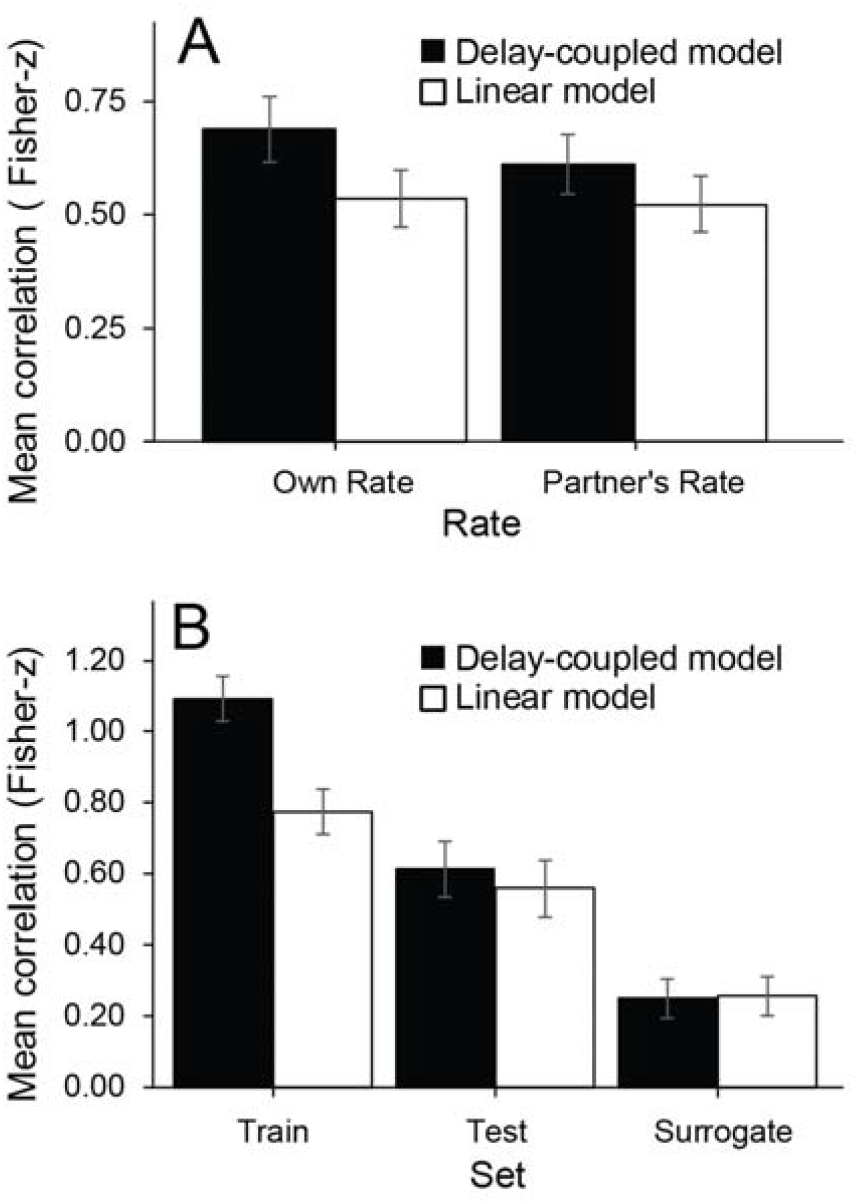
Correlations of observed asynchronies with model predictions: significant interactions. A: Rate × Model interaction. B: Model × Set interaction.

### Tests of estimated model parameters

A two-way ANOVA conducted on each of the estimated model parameters (κ and *ω*_*diff*_ values) by Cued Rate and Task did not reveal significant main effects or interactions. Individual differences in the model parameter values were investigated with a mixed model that included Musicianship, Task (Solo / Joint) and Faster / Slower Cued Rates as fixed effects. Table 1 shows the mean fitted κ values and *ω*_*diff*_ for musicians and nonmusicians at the Faster or Slower Rate and in Solo and Joint conditions. A significant main effect of Musicianship on the κ values, shown in Table 1, indicated that musicians had higher coupling values than nonmusicians, *F*(1, 17.11) = 5.62, *p* = .030; no other main effects or interactions reached significance in intrinsic (*ω*_*diff*_) and coupling (κ) parameters.

**Table 1.** Mean model parameter values for Musicians and Nonmusicians by Faster / Slower Cued Rate and Task

| | Musician $\kappa$<br>(SD) | Nonmus $\kappa$<br>(SD) | Mean $\omega_{diff}$<br>(SD) |
| --- | --- | --- | --- |
| Cued rate = Slower SPR |  |  |  |
| Solo | 4.89 (6.25) | 1.81 (3.32) | 71.33 (107.04) |
| Joint | 3.90 (4.69) | 2.02 (2.05) | 84.03 (83.80) |
| Cued rate = Faster SPR |  |  |  |
| Solo | 5.78 (8.06) | 1.70 (3.00) | 43.68 (52.04) |
| Joint | 6.73 (10.56) | 3.25 (3.74) | 58.18 (88.51) |

A significant positive correlation was observed between the intrinsic frequency (*ω*_*diff*_) and coupling (κ) parameters, both in Solo, *r*(40) = .45, *p* < .01, and in Joint tasks, *r*(40) = .47, *p* < .01. Thus, the analyses of individual differences were repeated for κ with *ω*_*diff*_ included as a covariate. The main effect of Musicianship was still observed, *F*(1, 17.12) = 7.60, *p* = .013, and a main effect of *ω*_*diff*_ was observed on κ, *F*(1, 147.70) = 121.12, *p* < .001. In addition, the Faster / Slower Cued Rate variable reached significance, *F*(1, 23.38) = 12.26, *p* = .01; partners showed greater coupling in response to the Faster cue (*M* = 3.21, *SD* = 4.58) than to the Slower cue (*M* = 4.43, *SD* = 7.37). There were no other significant effects or interactions.

### Social Interaction Ratings

Table 2 shows the median values for all participants on the social interaction questions. Participants in the Uncoupled pairs rated Pleasantness, Relationship and Synchronization as lower than participants in the Coupled pairs. Significant correlations were observed between participants’ ratings of pleasantness and relationship, *r(*18) = .65, *p* < .01, and between pleasantness and synchronization, *r*(18) = .75, *p* < .01. No other correlations reached significance between the social interaction questions, the mean and SD of asynchronies, and model parameters in the Joint condition.

**Table 2.**
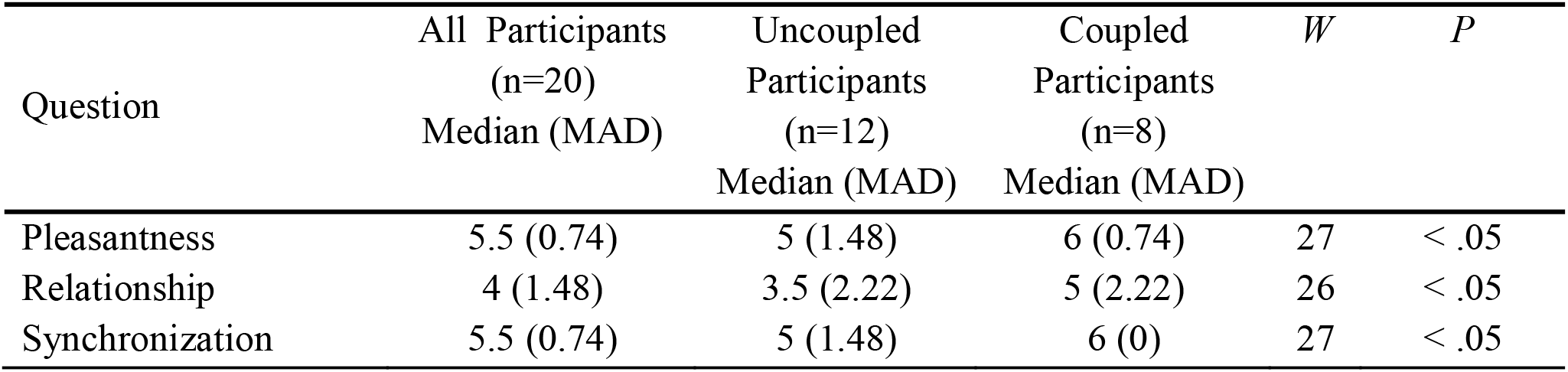
Median (and Median Absolute Deviation) ratings for social interaction questions by Coupled (delay-coupled model fit) and Uncoupled (linear model fit) pair members, with Wilcoxon tests (one-tailed).

| Question | All Participants<br>(n=20)<br>Median (MAD) | Uncoupled<br>Participants<br>(n=12)<br>Median (MAD) | Coupled<br>Participants<br>(n=8)<br>Median (MAD) | *W* | *P* |
| --- | --- | --- | --- | --- | --- |
| Pleasantness | 5.5 (0.74) | 5 (1.48) | 6 (0.74) | 27 | < .05 |
| Relationship | 4 (1.48) | 3.5 (2.22) | 5 (2.22) | 26 | < .05 |
| Synchronization | 5.5 (0.74) | 5 (1.48) | 6 (0) | 27 | < .05 |

Table 2 also shows the median values for the pairs whose model fits to asynchronies indicated Coupling, based on a better fit of the delay-coupled model (*n* = 6 pairs or 12 partners), or Uncoupling, based on a better fit of the linear model (*n* = 4 pairs or 8 partners). Both members of a pair were defined as “Uncoupled” if one or more partners in the pair failed to show Coupling (higher RMSE values for their asynchronies fit by delay-coupling model than by the linear model). As indicated in Table 2, Coupled pairs rated their partners significantly higher on the social variables of Pleasantness, Relationship and Synchronization than did the Uncoupled pairs. Thus, participants’ judgments of social interaction were greater for pairs in which the two partners displayed coupled behavior in their model fits to the observed asynchronies.

## Discussion

We compared musicians’ and nonmusicians’ synchronization accuracy in a tapping task, under social (joint) turn-taking conditions and under similar individual (solo) turn-taking conditions. Partners’ perceptions of social interaction were measured after their joint performances. A delay-coupling model and a linear model were applied to participants’ synchronization measures to capture their ability to couple with a regular auditory cue in the presence or absence of their partner. Both musically trained and untrained participants’spontaneous production rates showed a wide range (305 – 838 ms ITI), indicating that some (randomly paired) partners had to adapt a lot in order to synchronize at their partner’s rate, and thus the computational models had to account for large rate differences captured by the intrinsic frequency parameter. The partners’ musical experience and the difference in their spontaneous rates influenced the success of their synchronization, their perception of social interactions with their partner, and the computational models’ fits. We discuss each of these findings below.

Partners were more synchronous in the Joint turn-taking task when the auditory cue was set to their own spontaneous rate than to their partner’s rate consistent with the notion that individuals are attracted to their intrinsic frequencies and may have more difficulty tapping at rates farther from their spontaneous rates (Scheurich, Zamm, & Palmer, 2018). This result suggests that hearing one’s partner attempt to synchronize while one taps at the partner’s rate can reduce one’s own synchronization accuracy. It is important to note that the mere presence of the partner was not sufficient to disrupt synchronization, as the difference in synchronization accuracy between Solo and Joint tasks was not observed when the auditory cue was set to the partner’s own spontaneous rate. These findings suggest that the influence of a partner in a turn-taking task, in which there is no simultaneous action with the partner, may not always be beneficial.

The social context of the Joint tapping task did not improve partners’ synchronization accuracy overall, relative to the similar Solo tapping task performed at the same rates. This finding replicates Zamm et al. (2021) who reported that participants struggled more to learn a rhythmic sequence in a turn-taking context than in an individual context. The presence of a partner performing the same task may hinder participants’ performance, especially when their behavior is discrepant with that of their partner. This hypothesis is supported by the current finding that asynchronies were larger when the cued rate was set to the partners’ spontaneous rate in the Joint condition, but not in the Solo condition. Although this interpretation is at odds with the evidence of children’s improved synchronization in a social interaction context than in a solo context (Kirschner & Tomasello, 2009), several distinctions between these tasks can account for the different outcomes; for example, the more naturalistic turn-taking behavior in the current Joint task compared two social “equals” as participants, as opposed to the experimenter-child dyad of the previous study. In addition, the current task introduced auditory feedback from a partner, which may act as a perturbation, especially if the cued rate differed from the partner’s SPR. Finally, the observed partner differences in asynchrony were driven more by the faster partner who could not slow down to match the rate of their slower partner. This finding supports the view that there are large individual differences in rate flexibility (Scheurich, Zamm, & Palmer, 2018) and suggests that individuals with faster intrinsic frequencies are less flexible than slower individuals.

Participants’ perceptions of social interaction increased when their synchronizations were coupled with the auditory cue, based on the nonlinear model fits. Participants gave higher ratings to the task’s pleasantness, their perceived relationship with the partner, and ability to synchronize relative to the partner, when the two partners exhibited greater coupling in the Joint synchronization task, compared with pairs in which at least one partner did not exhibit coupling (that is, the linear model fit to asynchronies was equal to or better than the delay-coupling model). These findings are consistent with the observations that social connection and positive feelings are increased when synchronization with a partner is successful (Hove & Risen, 2009; Lumsden et al., 2014). The social interaction effects were modest in size, perhaps because each partner took turns synchronizing with a metronome, rather than synchronizing simultaneously (Hove & Risen, 2009; Lumsden et al., 2014).

The partners’ asynchronies were modeled with a delay-coupled model and with a linear model that contained only an intrinsic frequency term; the delay-coupled model fit better overall to the synchronization measures, for both musician and nonmusician pairs. Furthermore, the nonlinear model’s coupling parameter values were greater for musically trained pairs than for untrained pairs. This finding held even when differences in the partners’ intrinsic frequencies were included in the model fits to asynchronies. Musicians’ synchronization was less variable than that of nonmusicians, a finding that echoes previous observations that musicians’ motor variability is reduced relative to nonmusicians in tapping tasks (Nguyen et al., 2022).

An important exception to these model fits were the “uncoupled” synchronization trials of a few participants; a linear model best captured their steady drift away from the metronome, as shown in Figure 2, which indicates they did not couple with the auditory cue. These uncoupled fits also affected the partners’ subjective ratings of social interaction; tappers whose asynchronies were best fit by the linear model gave lower ratings of pleasantness, synchrony, and perceived relationship with their partner than did the pairs better fit by the delay-coupling model. Although most behavioral studies of synchronization report occasional participants whose synchronization measures do not indicate adjustment to an auditory cue, this is the first study to identify a model parameter that captures the distinction between coupled and uncoupled participants and to unite those differences with perceived social interaction measures.

These findings emphasize the importance of coupling strength in social interactions such as jointly performed auditory synchronization (Dotov et al., 2019). Each individual’s coupling strength increased when the difference between the cued rate and the model’s estimate of their intrinsic frequency increased. This is expected because a system’s coupling strength must increase when a large difference exists between the system’s intrinsic frequency and the external cued rate, in order to maintain synchrony. Thus, the delay-coupled model, by design, handles intrinsic rate differences between performers well. In contrast, weak anticipation models such as the linear phase correction model of Elliott et al. (2014) capture the timing of events using predictions from an internal timekeeper combined with an error correction parameter. Although that model was able to capture synchronization behavior during string quartet performances, it did not account for perceived social interaction among performers or their relationship with individual differences. Thus, the current findings suggest that the delay-coupling model is better suited to capture the wide range of individual differences in both production rates and synchronization, while offering an explanation for how perceptions of social interaction arise.

This study investigated turn-taking between partners in a novel task in which participants took turns synchronizing with a regular auditory cue. By presenting each partner’s spontaneous rate as auditory cues in both Solo and Joint conditions, the novel paradigm permitted us to evaluate the effects of intrinsic frequencies independent of the social effects of working with a partner. The influence of social interaction on synchronziation behaviors may differ when participants take turns in a less constrained interactive task (such as conversational speech). Future avenues of investigation include tapping performance in turn-taking tasks without an auditory pacing cue, and in tasks that permit overlap between participants’ performances at turn-switching points, similar to natural speech conversation. Paradigms that attempt to perturb turn-taking by means of adding more partners or increasing working memory demands (Maes et al, 2015) are also needed to confirm the relationship between temporal accuracy and social interaction.

In conclusion, turn-taking in a social context depends on who the partner is. When each partner synchronized their tapping with a regular auditory cue, the tappers were less accurate when they took turns with a partner than when they performed the same task individually. The amount of disruption in turn-taking increased with the difference in the partners’ spontaneous rates, consistent with dynamical systems that predict greater coupling necessary between oscillators operating at similar frequencies. A delay-coupled model accounted for participants’ synchronization through increased coupling values as the partners’ difference in spontaneous rates increased. Finally, individuals with greater coupling values perceived the interaction as more pleasant, more synchronous, and had a stronger relationship with their partner than individuals with smaller coupling values. Future studies may address how individual differences in synchronization performance affect social interaction. The finding that some individuals make better partners even when they take turns (do not synchronize simultaneously) suggests intriguingly that perceived social interaction arises in part from the perceived degree of coupling between individuals with external auditory events in the world.

## Notes

This research was supported by an NSERC-CREATE Postdoctoral Award to V. Bégel and by NSERC Grant 298173 and a Canada Research Chair to C. Palmer. We thank Jocelyne Chan, Sasha Sorger Brock, and Vivian Qiang for their assistance.

